# Effects of nicotinic receptor antagonism on nicotine and THC self-administration in a model of polysubstance use

**DOI:** 10.1101/2025.09.05.674477

**Authors:** Mary M. Torregrossa, Tamara Racic, Samantha L. Baglot, Sierra J. Stringfield, Matthew N. Hill, Alan Sved

**Affiliations:** Department of Psychiatry, University of Pittsburgh, Pittsburgh, PA 15219; Department of Neuroscience, University of Pittsburgh, Pittsburgh, PA 15213; Departments of Cell Biology and Anatomy & Psychiatry, Cumming School of Medicine, Hotchkiss Brain Institute and The Mathison Centre for Mental Health Research and Education, University of Calgary, Calgary, Canada

**Author notes:** Corresponding Author: Mary M. Torregrossa 450 Technology Drive, Bridgeside Point II, Room 228 Pittsburgh, PA 15219.

**Keywords:** Nicotine, THC, polysubstance use, self-administration, nAchR antagonist

## Abstract

**Background:** Most substance users are polysubstance users; however, little is known about how the combined use of different drugs affects the course of substance use disorder or effectiveness of treatment. Notably, co-use of cannabis and nicotine is very common, and we previously demonstrated that nicotine enhances the self-administration of a synthetic cannabinoid receptor agonist and the primary psychoactive phytocannabinoid in cannabis, Δ-9-tetrahydrocannabinol (THC).

**Methods:** Here we aimed to further investigate the patterns of nicotine and THC self-administration when available in a concurrent choice model, and to determine the effects of nicotinic acetylcholine receptor (nAchR) antagonists on nicotine-enhanced THC self-administration in male and female rats.

**Results:** During concurrent choice, nicotine availability increased THC self-administration in females without affecting THC metabolism, while THC availability decreased nicotine self-administration and preference in females relative to when nicotine and saline were concurrently available. In females, THC self-administration was reduced by the β2/4 subunit-containing nAchR antagonist dihydro-beta-erythroidine (DHβE) in both nicotine and saline concurrent availability groups; while nicotine self-administration was reduced in both sexes by the α7nAchR antagonist methylylcaconitine (MLA), but only in rats that had concurrent access to THC. The nonspecific nAchR antagonist mecamylamine had minimal effects in the concurrent choice model, but it prevented nicotine-induced enhancement of THC self-administration when nicotine was given prior to a single choice THC only self-administration session.

**Conclusions:** Thus, behavioral regulation of self-administration is differentially influenced by nAchR subtypes depending on the availability of other substances, which has implications for the efficacy of treatments in the context of polysubstance use.

**Significance Statement:** Polysubstance use is extremely common, but very understudied in both clinical and preclinical research. The results presented here highlight that pharmacological modulators of drug reinforcement can differ when multiple drugs are available simultaneously, highlighting the importance of investigating potential treatments for substance use disorders in the context of polysubstance use.

## Introduction

Polysubstance use (PSU) is extremely common, with the majority of substance use treatment seekers reporting the use of multiple substances ^1^. Not only is the prevalence of PSU high, but the consequences of PSU are also thought to be worse than for individuals who use single substances. There is evidence for worse psychosocial and cognitive disruptions in those that use multiple substances, and this is particularly true in youth ^1–5^. PSU is also associated with increased consumption of both substances, which can lead to worse physical health outcomes ^5,6^. Finally, people with PSU often find it harder to quit either substance, and the continued use of one substance can lead to relapse to the use of the other substance.

One of the most common substances to be used with any other drug of abuse is nicotine. The majority of psychostimulant, opioid, and cannabis users also report the use of nicotine ^3^. Cannabis and nicotine co-use is particularly common ^7^, with higher overall incidence of cannabis use in the general population and up to 80% of cannabis users reporting that they also use nicotine ^8^. In addition to using both drugs on separate occasions, nicotine and cannabis products are commonly combined so that they are consumed simultaneously, and nicotine is often reported as a “chaser” to cannabis use ^2,8,9^. Recent epidemiological studies indicate that the co-use of nicotine with THC containing products is increasing. Those that use both drugs at the same time (whether delivered at the same time or right after each other) are reported to have worse psychosocial and functional outcomes than those individuals that consume both, but on separate occasions ^2^.

To increase our understanding of nicotine/cannabis PSU we previously investigated the effects of pre-session nicotine administration on the self-administration of both the synthetic cannabinoid receptor agonist WIN 55,212-2 (WIN) and the primary psychoactive component of cannabis, Δ9-tetrahydrocannabinol (THC). We also developed a model of dual IV catheter concurrent choice self-administration of nicotine and WIN. In all cases, nicotine administration or concurrent nicotine availability increased self-administration of both cannabinoids. Thus, nicotine appears to produce reinforcement-enhancing effects for cannabinoids, similar to prior research demonstrating that nicotine enhances responding for other mildly reinforcing stimuli ^10–13^. Prior research on nicotine’s reinforcement enhancing effects has found that this is mediated through activation of β2 subunit containing nicotinic acetylcholine receptors (nAchRs), but not α7nAchRs ^14–17^. However, this may not be the case for cannabinoids as THC self-administration in squirrel monkeys was previously shown to be attenuated by kynurenic acid mediated negative allosteric modulation of α7nAchRs ^18,19^. Therefore, in the present study we investigated the role of different nAchR subtypes on concurrent IV self-administration of nicotine and THC in male and female rats, examining both overall effects on self-administration and relative preference between each substance and its respective vehicle. By investigating how nAchR antagonists alter reinforcing properties of nicotine and THC as well as choice preference between them, this study may help identify novel therapeutic targets for interventions aimed at nicotine and cannabis dependence or polysubstance use.

## Materials and Methods

### Animals

Adult male and female Sprague Dawley rats purchased from Envigo (Indianapolis, IN) weighed approximately 250 g and 225 g, respectively, on arrival. Rats were single-housed on a 12-12 hr reverse light cycle in a temperature and humidity-controlled room. Food and water were provided *ad libitum*. All animal procedures were conducted in accordance with the guidelines of the National Institutes of Health *Guide for the care and use of Laboratory Animals* and were approved by the University of Pittsburgh Institutional Animal Care and Use Committee.

### Drugs

Nicotine hydrogen tartrate salt (MP Pharmaceuticals, Solon, OH) was dissolved in 0.9% sterile saline. All nicotine doses are expressed as free base. Δ-9-tetrahydrocannabinol (THC, generously provided by the National Institute on Drug Abuse’s Drug Supply program) was suspended in 100–200 μl of Tween 80 and the ethanol solvent was evaporated using nitrogen gas, as described previously ^20^. The selective α7nAchR antagonist, methylylcaconitine citrate (MLA, Sigm-Aldrich) was dissolved in saline and given at a dose of 10 mg/kg, i.p. The β2/4 subunit containing nAchR antagonist, dihydro-beta-erythroidine hydrobromide (DHβE, ApexBio) was dissolved in saline and given at a dose of 3 mg/kg, s.c. The non-specific nAchR antagonist mecamylamine hydrochloride (MEC, Sigma-Aldrich) was dissolved in saline and given at a dose of 1 mg/kg, i.p. Doses were chosen based on effective doses in the literature, including our own publications ^14–16^.

### Surgical Procedures

Catheters were constructed and implanted either into one or both jugular veins as described previously ^21^. Rats were allowed to recover for at least 5 days and catheters were flushed daily with a 0.1 ml solution containing 30 U/ml heparin and 66.67 mg/ml gentamicin. Catheter patency was confirmed periodically with methohexital (5 mg/kg) and after the final self-administration session.

### Apparatus

Self-administration sessions occurred in standard operant chambers (Med-Associates, St Albans, VT) containing two nose poke ports with white stimulus lights located directly above each port. A red houselight was located on the same wall. Syringe pumps were connected to an infusion swivel with tubing to connect to catheter ports. During dual self-administration experiments, two separate syringe pumps were connected to a dual channel fluid swivel to allow for access to two distinct solutions during each session, as previously described ^21^.

### Intravenous Self-Administration Sessions

Rats completed daily 1-hour self-administration sessions. Rats in experiments self-administering one drug solution were randomly assigned to active and inactive nose poke ports, and responses into the inactive port were recorded but did not have programmed consequences. Responding in the active port resulted in the intravenous delivery of drug solution and 1 s illumination of the stimulus light located directly above the port, followed by a mandatory 60 s timeout where responses into the active port were recorded but did not result in drug infusion. For dual self-administration sessions, both nose poke ports were active and resulted in an infusion of a specific solution, followed by the timeout period. Responding in one port resulted in an infusion of the associated drug along with presentation of the stimulus light located directly above the port. Infusion times were adjusted based on body weight to administer the specified drug dose.

#### Experiment 1. Effect of nAchRs Antagonists on Dual-Catheter Concurrent Choice Self-Administration

Male and female rats were implanted with dual catheters to allow for concurrent access to two separate drug solutions during a single self-administration session. Rats were trained to associate one nosepoke port with administration of nicotine (NIC, 30 µg/kg/infusion), THC (escalating doses), or saline (SAL), and the other nosepoke port with administration of an alternative solution. When rats were assigned to a THC paired nosepoke, THC was given at a dose of 3 µg/kg/infusion for the first 3 days, 10 µg/kg/infusion for the next 4 days, and were then switched to a unit dose of 30 µg/kg/infusion for the remaining days of self-administration . Rats were randomly assigned to groups where they had access to pairs of drug solutions: THC and NIC (males N=23, females N=18), THC and SAL (males N=20, females N=21), NIC and SAL (males N=27, females N=14). Both solutions were available throughout the 1-hour session. Rats self-administered for a minimum of 18 days on an FR1 schedule of reinforcement prior to antagonist administration testing.

During the antagonist testing phase, rats were randomized to receive pre-session injections of the vehicle solution or one of the three antagonists across multiple testing days. Vehicle, MEC, and DHβE were given 10 minutes, and MLA 15 minutes prior to self-administration sessions in a randomized order within and between groups of rats. All rats received a vehicle test day for each antagonist tested, and some combination of the antagonists, but not in the same order, and not all rats received all three antagonists. In between test days, rats underwent at least 1 concurrent choice session without treatment before testing again with another agent. Catheter patency was also confirmed periodically, and rats were removed from the study at the point of catheter failure. The final group sizes for each treatment are as shown in Table 1.

**Table 1.**

| **TABLE 1.** |  |  |  |  |
| --- | --- | --- | --- | --- |
| **NUMBER OF ANIMALS IN EACH TREATMENT GROUP** |  |  |  |  |
| **CONCURRENT ACCESS GROUP** | **ANTAGONIST TREATMENT** |  |  |  |
|  | **VEHICLE** | **DHBE** | **MLA** | **MEC** |
| **THC+NIC FEMALE** | **13** | **13** | **9** | **5** |
| **THC+NIC MALE** | **14** | **17** | **8** | **8** |
| **THC+SAL FEMALE** | **10** | **8** | **6** | **6** |
| **THC+SAL MALE** | **13** | **13** | **8** | **7** |
| **NIC+SAL FEMALE** | **12** | **13** | **10** | **4** |
| **NIC+SAL MALE** | **18** | **13** | **10** | **10** |

#### Experiment 2. Effect of nAchR Antagonists on Nicotine-Enhanced THC Self-Administration

Male and female adult rats were implanted with single lumen catheters and were trained to self-administer escalating doses of THC in a traditional single-choice paradigm. Rats received non-contingent pre-session injections of either nicotine (0.3 mg/kg) or saline as previously published ^21^ (nicotine: N=10 female, N=8 male; saline: N=8 female, N=7 male). Rats self-administered 3 µg/kg/infusion THC for days 1-3 of training, followed by 10 µg/kg/infusion on days 4-7, and 30 µg/kg/infusion beginning on day 8 and continuing until the end of the experiment. Rats self-administered for at least 19 days prior to antagonist testing. Rats were randomly assigned to antagonist and vehicle testing in a counterbalanced order across days, though not all rats received all three antagonists, depending on catheter patency. Rats received injections of the antagonist or vehicle 10 or 15 minutes (as detailed above) prior to pre-session injection of nicotine or saline, which occurred 10 minutes prior to the start of the self-administration session. Rats received at least one untreated self-administration session between antagonist tests, maintaining their saline or nicotine pre-session treatment.

#### Experiment 3. Effect of Nicotine on THC and Metabolite Concentrations

In order to determine if the administration of nicotine might influence THC metabolism in a manner that might explain why nicotine increases THC self-administration, an additional experiment was completed similar to Experiment 2, where pre-session nicotine injections were given during single choice THC self-administration (nicotine: N=7 female, N=6 male (1 female and 1 male brain sample were not usable); saline: N=6 female, N=6 male). Immediately following the final self-administration session, rats were euthanized, and trunk bloods and brains were collected for analysis of plasma and brain levels of THC and metabolites.

### Determination of Tissue Levels of THC and Metabolites

Blood and tissue samples were collected immediately after the 1-hour self-administration session described above. Rats were euthanized by rapid decapitation and at least 500 uL trunk blood was collected in heparin coated tubes while brains were collected, frozen on dry ice, and stored at -80 °C. Blood samples were centrifuged at 1500 × g for 10 minutes at 4°C before the plasma was collected and stored at -20 °C. Plasma and brain samples were then shipped on dry ice for analysis of THC and its metabolites using an established protocol ^22^. Deuterated (d3) standard of THC, 11-OH-THC, and carboxy-THC (THC-COOH) was dissolved in <u>acetonitrile</u> (ACN) to a final concentration of 1.0 mg/ml. An internal standard was prepared with 1:2 methanol water to contain 10 ng/ml of each compound. Glass tubes containing 2 ml of ACN and 100 µl of internal standard were prepared for plasma and brain samples. Plasma samples were thawed to room temperature and 100 µl of plasma were added to each tube. Brain samples were kept frozen on dry ice before being weighed and placed in the glass tube for homogenization using a glass rod. All samples were then sonicated for 30 min in an ice bath and then stored at −20°C overnight to precipitate proteins. The next day, samples were centrifuged briefly (3–4 min at 4°C) at 1800 RPM to separate particles from the supernatant. The supernatant was transferred to a new glass tube and placed under nitrogen gas until almost evaporated. Glass tubes were washed with 250 µl of ACN and then left to evaporate fully. Following evaporation, all samples were resuspended using 100 µl of 1:1 methanol deionized water. Samples underwent two rounds of centrifugation at 15000 RPM for 20 min at 4°C. Resuspended samples were stored at −80°C until analysis using LC-MS/M by the Southern Alberta Mass Spectrometry (SAMs) facility. Values were normalized to tissue weight or plasma volume and converted to ng/g or ml, respectively, for statistical analysis and graphing.

### Data Analysis

Statistical analysis was completed using GraphPad Prism version 10.4.1 (GraphPad Software, San Diego, CA). Drug intake for both THC and NIC was compared across self-administration sessions by repeated measures 3-way ANOVA or mixed effects analysis when there were missing values including factors of self-administration group (alternative option in concurrent experiments, or pre-treatment group in single access experiments), sex, and antagonist pre-treatment when appropriate. Significant effects were further analyzed by two-way or one-way ANOVA and further Tukey’s post-hoc tests, when applicable. Geisser-Greenhouse correction was applied when the assumption of sphericity in the data was violated. Chi-square analysis was performed to determine differences in proportions of rats preferring THC or NIC, and simple regressions were performed to determine relationships between THC infusions self-administered and brain and plasm levels. α = 0.05 was used for all analyses.

## Results

### Effects of nicotine-THC concurrent availability on self-administration patterns

Our previous work investigated the effect of concurrent nicotine-WIN availability on WIN self-administration, but we had not previously determined if concurrent nicotine would have the same effect on THC self-administration. Thus, we first determined nicotine and THC self-administration patterns under concurrent choice. THC self-administration was compared in males and females that either had nicotine or saline concurrently available. Three-way repeated measures (rm) mixed effects analysis of the number of THC infusions earned across days found significant effects of day and sex, and interactions between nicotine versus saline access with day and sex. Follow-up analyses found sex-specific effects of nicotine on THC self-administration over multiple days. In females, concurrent access to nicotine significantly increased THC infusions. Males with concurrent access to saline or nicotine exhibited greater THC self-administration than females in the THC+saline group, but nicotine availability did not affect male THC intake (Fig. 1a). Nicotine self-administration analyzed across all days of self-administration was not significantly different based on sex or concurrent access group (Fig. 1b). However, given that the most prominent differences in behavior emerged in the latter days of self-administration, we compared the average of the last 3 days of self-administration between groups. For THC self-administration, 2-way ANOVA revealed a significant effect of concurrent access to nicotine versus saline (F_(1,78)_ = 6.16, p=0.015) and a significant interaction between sex and solution (F_(1,78)_ = 7.27, p=0.009). Tukey’s post-hoc tests found that females with concurrent access to nicotine self-administered significantly more THC than those in the saline group (p=0.003), and that in rats with concurrent access to saline, males self-administered more THC than females (p=0.038; Fig. 1c). Nicotine self-administration, on the other hand, was reduced in females with concurrent access to THC (main effect of concurrent THC access (F_(1,78)_ = 7.67, p=0.007); effect driven by difference between females, p=0.03), while no other significant differences were detected (Fig. 1d).

**Figure 1.**
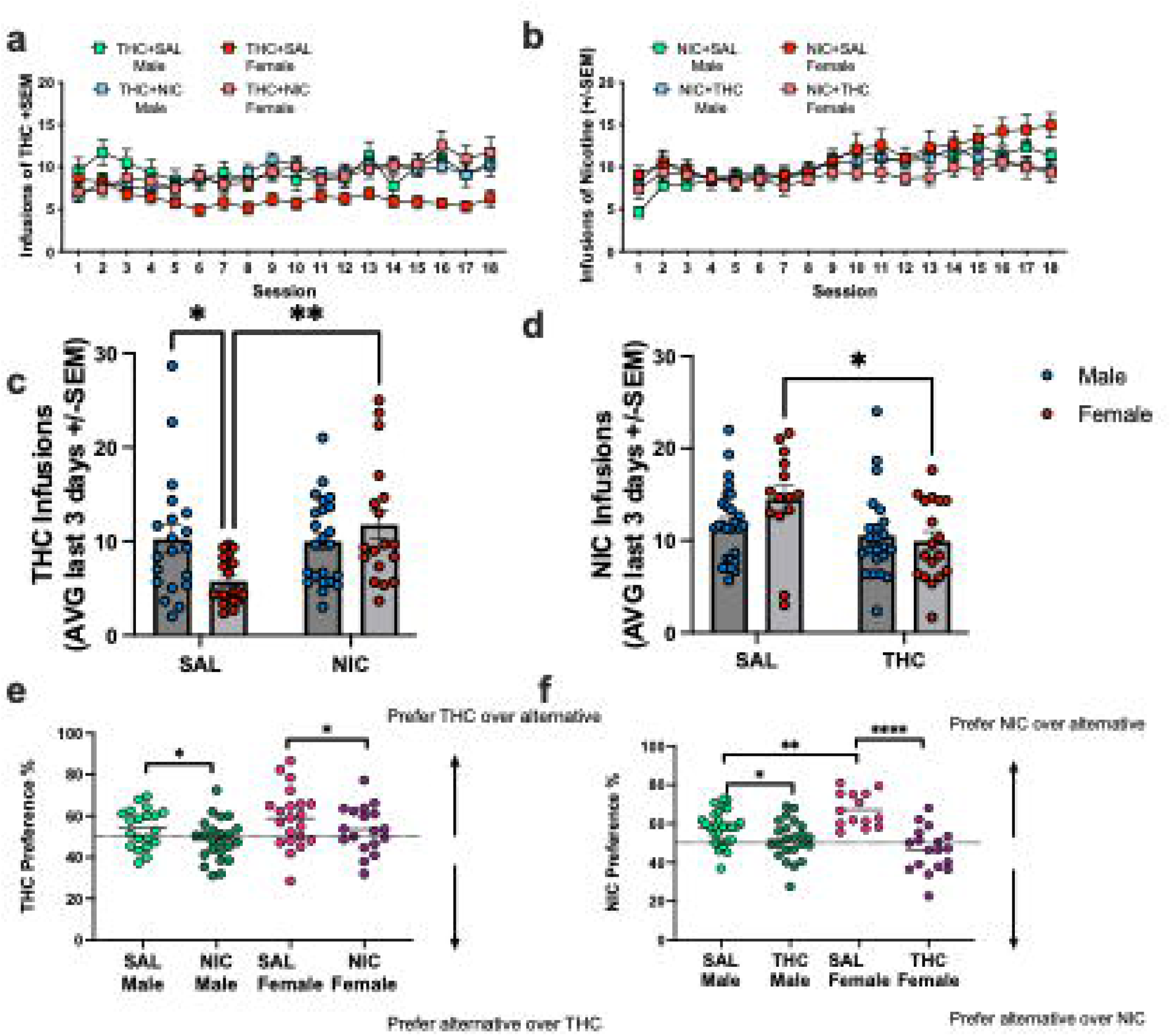
Concurrent access self-administration of nicotine, THC, and/or saline in male and female rats. a) Mean infusions of THC earned per day in rats self-administering THC where either saline (SAL) or nicotine (NIC) is concurrently available. b) Mean infusions of NIC earned per day in rats self-administering NIC where either saline (SAL) THC is concurrently available. c) Average THC infusions earned on the last 3-days of self-administration. Females with concurrent access to NIC self-administered more THC than females with concurrent access to SAL. No differences were observed in males. d) Average NIC infusions earned on the last 3-days of self-administration. Females with concurrent access to THC self-administered less NIC than females with concurrent access to SAL. No differences were observed in males. e) Preference scores for THC averaged over the last 3 days of self-administration. Concurrent access to NIC reduced THC preference. f) Preference scores for NIC averaged over the last 3 days of self-administration. Concurrent access to THC reduced NIC preference. Infusions earned are presented as mean ± SEM, * p<0.05, ** p<0.01, ****p<0.0001.

Given that concurrent access to THC and nicotine appeared to shift self-administration patterns relative to saline availability, particularly in females, we next examined the differences in preference for each of the two solutions by calculating the percentage of responses directed to the THC or NIC paired nose poke averaged over the last 3 days of self-administration. Preference for the THC paired nose poke was significantly attenuated by concurrent access to nicotine in both sexes (Two-way ANOVA main effect of solution (F_(1,78)_ = 4.95, p=0.029); Fig. 1e). Similarly, nicotine preference was significantly attenuated by concurrent access to THC, while females showed greater preference for nicotine over saline relative to males (Two-way ANOVA interaction between sex and solution (F_(1,78)_ = 10.6, p=0.002) followed by Tukey’s post-hoc tests; Fig. 1f). Overall, these findings suggest that rats direct more behavior to a THC or nicotine paired nose poke when the alternative option is a saline infusion, However, when both THC and nicotine are concurrently available, rats tend to allocate their behavior to both options roughly equivalently, though a range of individual differences are observed between which solution is preferred.

### Effects of nicotinic receptor antagonists on self-administration patterns during nicotine-THC concurrent availability

Following the 18 days of training on concurrent choice self-administration, we began testing the effects of nicotinic receptor antagonists on THC and nicotine self-administration patterns. Rats were given injections of the selective α7nAchR antagonist, MLA, the β2/4 subunit containing nAchR antagonist, DHβE, the non-specific nAchR antagonist, mecamylamine, or vehicle in a randomized order across days of self-administration, with drug-free self-administration days intermixed with antagonist treatments. Using three-way rm mixed effects analysis we compared differences in THC infusions earned based on sex, concurrently available solution, and antagonist treatment. We found a significant effect of solution (F_(1,46)_ = 4.93, p=0.031), an antagonist x solution interaction (F_(3,80)_ = 2.91, p=0.039), and a trend toward an interaction between solution and sex (F_(1,46)_ = 3.32, p=0.075). The main effect of solution appears to be primarily driven by the tendency for overall lower THC self-administration in the females with concurrent saline availability. Further mixed effects analysis of the effects of solution and antagonist revealed a significant antagonist effect (F_(2.33,27.2)_ = 4.75, p=0.013), but no interaction with solution (F_(3,35)_ = 0.58, p=0.63). Further analysis of antagonist effects within sex, found that DHβE pretreatment significantly reduced THC infusions earned relative to vehicle, selectively in females, regardless of which solution was concurrently available (p=0.004; Fig. 2a).

**Figure 2.**
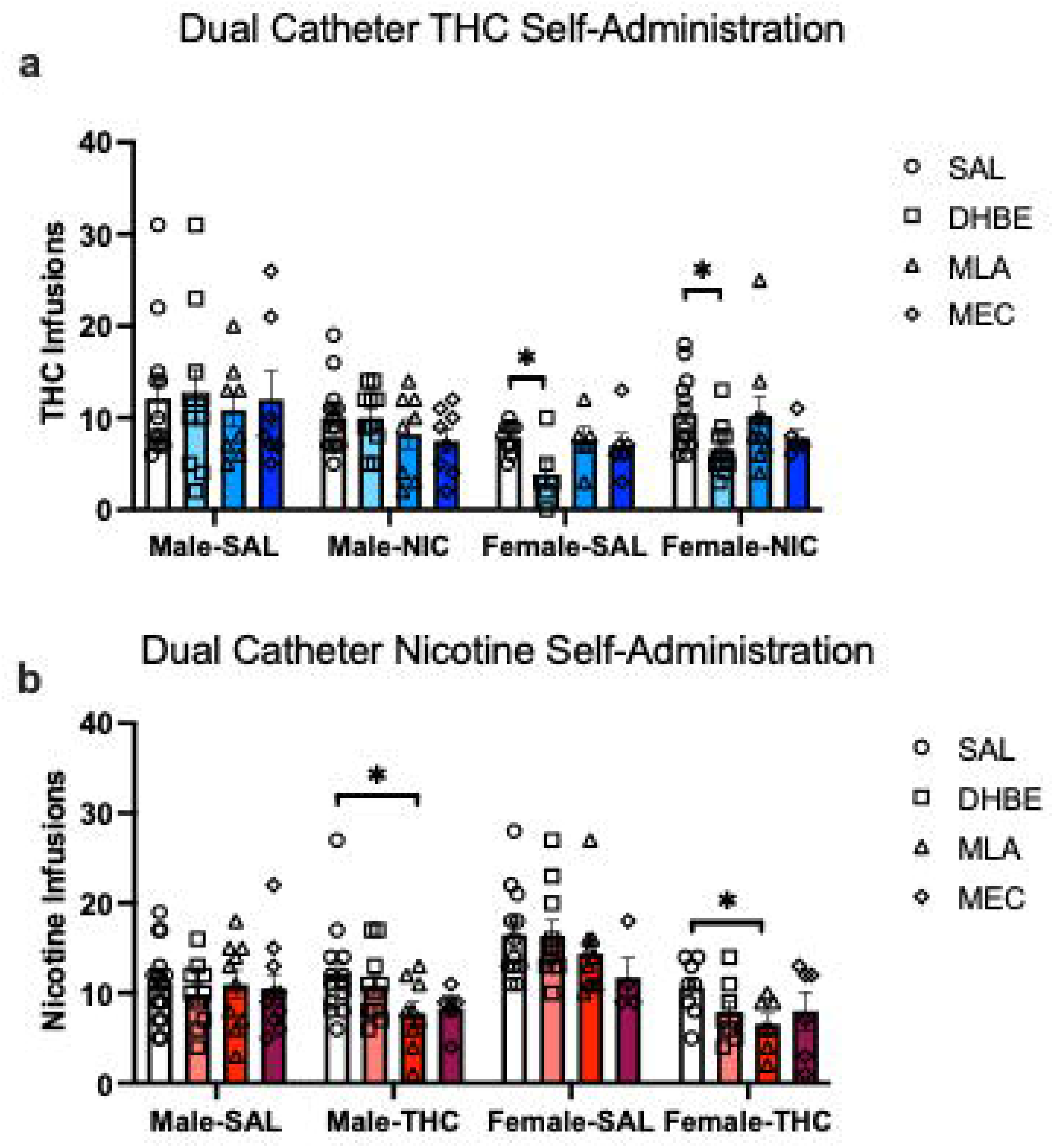
Effects of nAchR antagonists during concurrent access self-administration. a) THC infusions earned in male and female rats with concurrent access to NIC or SAL on single session test days where rats received a pre-session of an antagonist or vehicle. Females show reduced THC self-administration after treatment with DHβE. b) NIC infusions earned in male and female rats with concurrent access to THC or SAL after treatment with nAchR antagonists or vehicle. Nicotine self-administration was reduced in male and female rats with concurrent access to THC. Infusions earned are presented as mean ± SEM, * p<0.05.

Similar analysis of nicotine infusions earned revealed a different pattern of results. We found main effects of antagonist (F_(2.47,68.4)_ = 4.65, p=0.008), solution (F_(1,52)_ = 16.0, p=0.0002), and a sex x solution interaction (F_(1,52)_ = 8.29, p=0.006). To determine what drove the effects of antagonist treatment, we performed follow-up ANOVAs and found a significant effect of MLA compared to saline treatment in both males and females (p=0.01) and a marginal effect of MEC (p=0.052), but only in rats that had concurrent access to THC. No significant effects of any nAchR antagonist treatment were found in the concurrent saline access groups (all p>0.3, Fig. 2b). In the THC+NIC group, nicotine self-administration appeared to be reduced to some degree by all three antagonists, and they do not differ from each other statistically.

Given the different patterns of nAchR antagonist effects based on whether THC or NIC is being self-administered and what solution is concurrently available, we were curious if nAchR antagonist treatments shifted preference for one solution to the other. We calculated the preference score for each animal on the day they received the vehicle pretreatment for each antagonist and their preference for each substance on the day of antagonist treatment. In this analysis, we classified any rat that allocated greater than 55% of its responses to the THC port as THC preferring, rats that allocated greater than 55% of their responses to the NIC port as NIC preferring, and rats that dedicated between 45-55% of their responses to each port as neutral, or not strongly preferring one the substance over the other. We then performed chi-squared analysis to compare the proportion of animals classified under each preference category after antagonist treatment. The three different vehicle treatment days did not significantly differ from each other, so we combined them to get an overall proportion of preferences under control conditions, and compared that proportion to each antagonist. MEC pretreatment led to a significant difference in the proportion of animals preferring THC and nicotine, with a marked decrease in rats choosing THC over nicotine, and an increase in neutral responding (Chi-square = 6.2, p=0.045). The MEC group was also significantly different from MLA (Chi-square = 8.09, p<0.018) and DHβE (Chi-square = 31.6, p<0.001) treatment days (Fig. 3). Therefore, it appears that broad spectrum antagonism of nAchRs either decreases the reinforcing value of THC and/or leads to an increase in nicotine seeking in otherwise THC-preferring animals in an attempt to obtain the expected effects of nicotine. Interestingly, preference for the nicotine port was not dramatically affected by nAchR antagonism.

**Figure 3.**
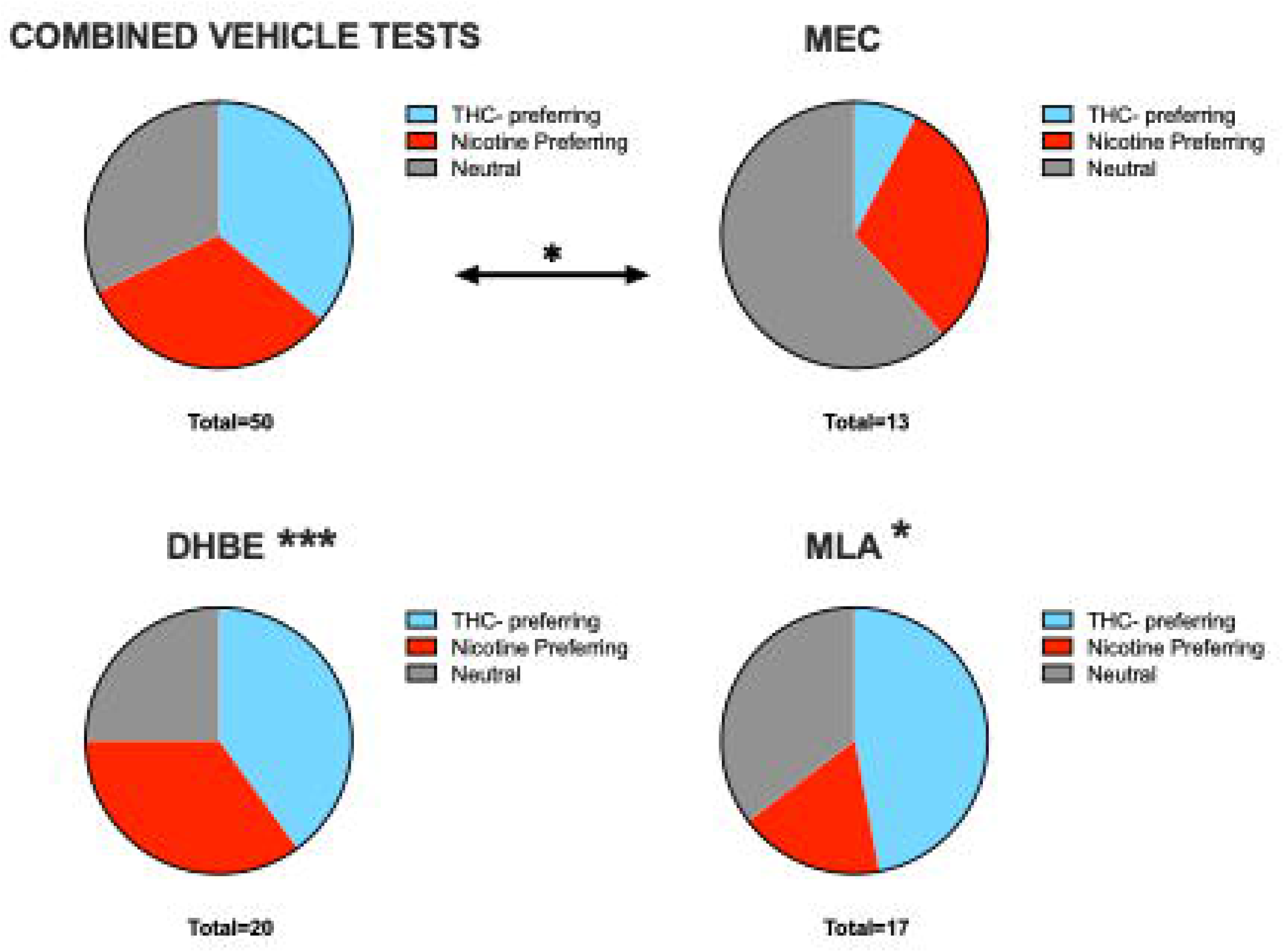
Effects of nAchR antagonists on the proportion of rats that prefer NIC versus THC. Rats were classified as preferring NIC or THC if the allocated more than 55% of their responses to the associated nose poke port. Rats between 45-55% preference were classified as neutral, non-preferring rats. MEC treatment reduced preference for THC relative to vehicle and to treatment with MLA or DHβE. * p<0.05, *** p<0.001.

### Effects of nicotinic receptor antagonists on nicotine-enhanced THC self-administration under single choice conditions

Overall, these data suggest that concurrent access to THC and NIC leads to differential effects of nAchR antagonism than what has been previously reported in the literature when only a single solution is available. Therefore, we wanted to compare the results from the concurrent access choice model to the single choice THC self-administration model, with or without pre-session experimenter-administration of nicotine. Thus, rats received either 0.3 mg/kg nicotine or saline 30 minutes prior to 1-hr THC single catheter self-administration sessions. We found that both male and female rats receiving daily nicotine pre-treatment developed an increase in the number of THC infusions earned during daily self-administration sessions relative to the saline pre-treatment group, replicating our previous findings ^21^ (Fig. 4a) [3way rmANOVA: main effect of day (F_(4.038,117.1)_ = 2.98, p=0.022); main effect of pre-session treatment (F_(1,29)_ = 13.64, p<0.001); day x pre-session treatment interaction F_(18,522)_ = 4.9, p<0.001); and no effect of sex F_(1,29)_ = 1.2, p=0.281) or interactions with sex (all p>0.27)].

**Figure 4.**
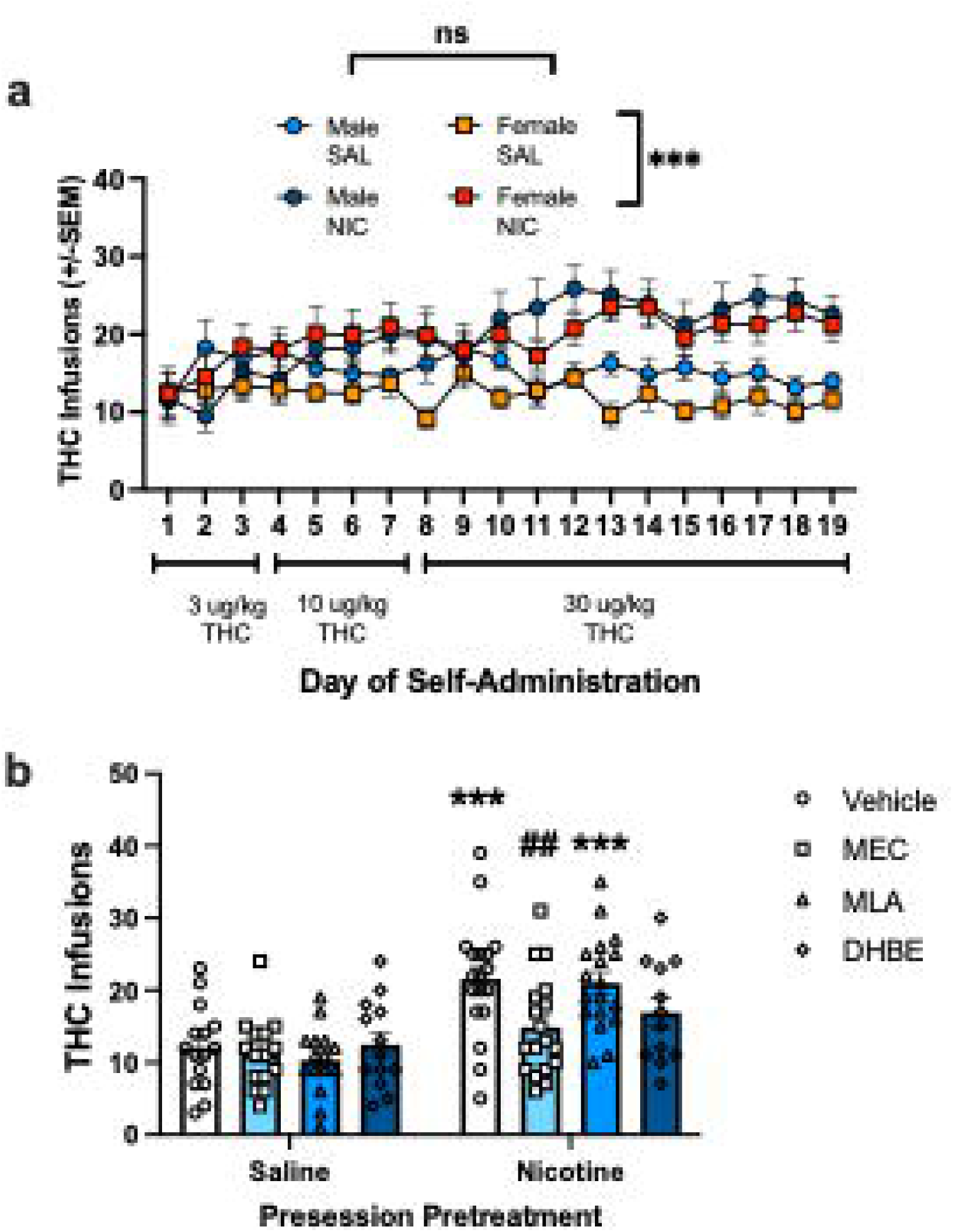
Effects of nAchR antagonists on the nicotine reinforcement enhancement effects for THC. a) Mean daily infusions earned in male and female rats with presession injection of NIC or SAL. Both male and females pre-treated with NIC earned more THC infusions during self-administration than the SAL pre-treatment group. b) THC infusions earned on discrete antagonist treatment test days with males and females combined. Nicotine’s reinforcement enhancing effect was significantly blocked by MEC and partially attenuated by DHβE. *** p<0.001 comparing NIC to SAL pre-treatment groups, ##p<0.01 comparing MEC to Vehicle pre-treatment.

Then, on testing days, rats received either a vehicle or nAchR antagonist treatment prior to pre-session injection of nicotine or saline in a pseudo-randomized manner. We combined males and females for analysis, as there were not baseline sex differences based on pre-treatment group. Mixed effects analysis comparing antagonist effects across pre-session treatment groups found significant effects of pre-session treatment (F_(1,31)_ = 16.82, p<0.001), antagonist (F_(2.587,73.29)_ = 3.2, p=0.035), and a pre-treatment by antagonist interaction (F_(3,85)_ = 3.99, p=0.01). Post-hoc analysis found no effects of any of the antagonists on THC self-administration in the saline pre-treatment group, consistent with the idea that nicotinic receptors are not required for THC reinforcement on its own. However, in the nicotine pre-treatment group, the enhancement of THC self-administration by nicotine was blocked by MEC (p=0.006), largely attenuated by DHβE (p=0.08), and not affected by MLA (p=0.73) (Fig. 4b). These findings are consistent with prior work identifying the receptor mechanisms mediating nicotine’s reinforcement enhancing effects in single choice settings ^10,12^.

### Assessment of THC bioavailability with nicotine pre-treatment

To determine if any of nicotine’s effects on THC reinforcement are due to differences in THC metabolism/bioavailability, we performed an additional experiment where rats received pre-session injections of nicotine or saline prior to single-choice THC self-administration sessions as above. Immediately following the 35th session, rats were euthanized, and trunk bloods and brains were taken for analysis of THC concentrations. First, we confirmed that nicotine pre-treatment increased THC self-administration by analyzing infusions earned in 15 min bins across the last 60 min session. As expected, both male and female rats pre-treated with nicotine obtained more THC infusions than rats pre-treated with saline, particularly in the last 45 min of the session (Fig. 5a,b). Analysis of cumulative infusions earned by 3-way rmANOVA found significant effects of bin (F_(1.22,25.61)_ = 41.7, p<0.001), pretreatment group (F_(1,21)_ = 6.89, p=0.016), and bin x pretreatment interaction (F_(3,63)_ = 7.28, p<0.001), but no effect of sex or interactions with sex (all p>0.25).

**Figure 5.**
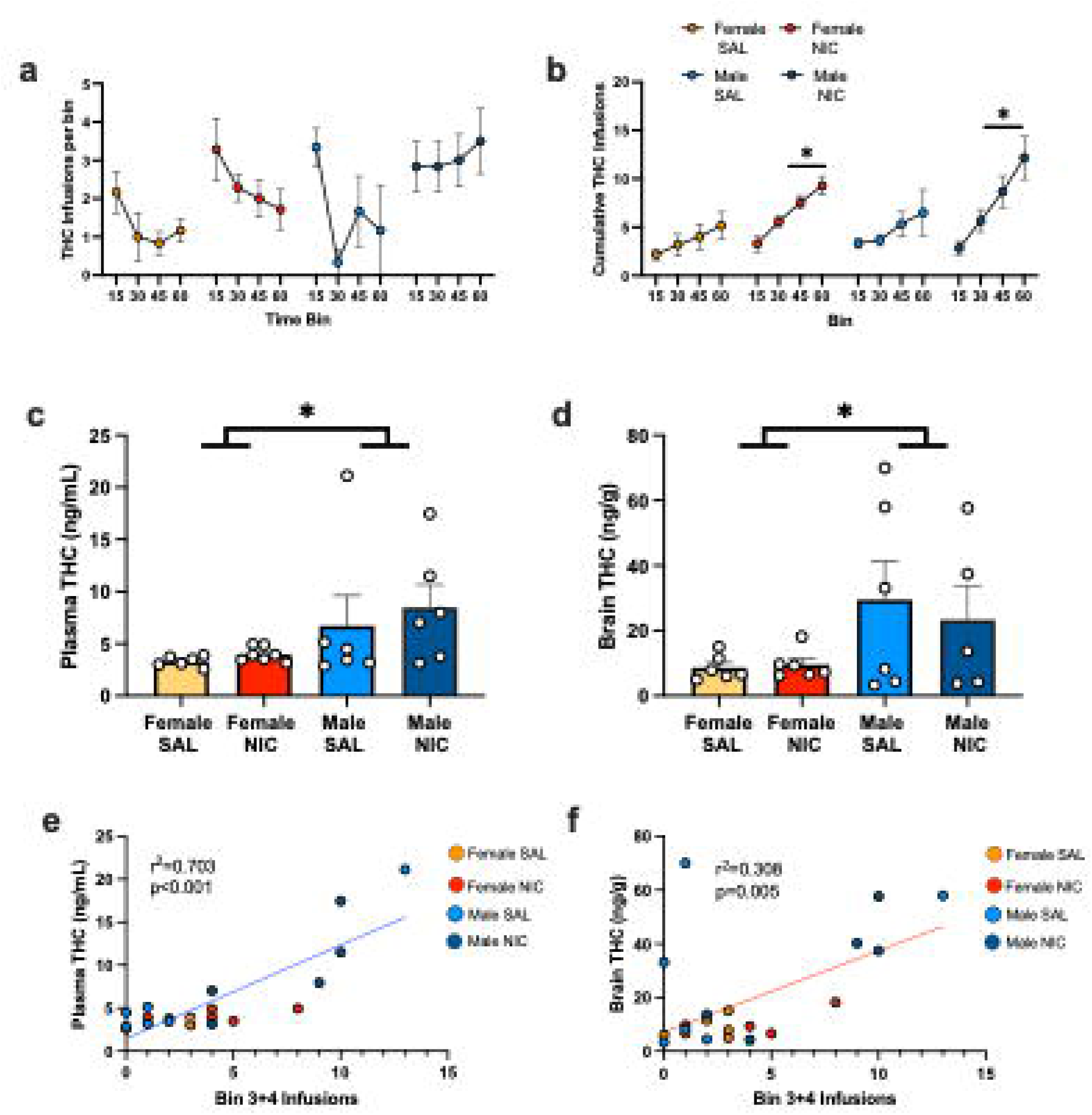
Effects of nicotine and sex on THC plasma and brain concentrations after self-administration. a) Mean THC infusions per 15 min bin during 60 min self-administration sessions in male and female rats pre-treated with SAL or NIC before the session. b) Cumulative THC infusions earned over 15 min bins of THC self-administration between groups. NIC pretreatment leads to a progressive increase in THC infusions earned in males and females. c) Plasma THC concentrations measured in rats after the self-administration session shown in a-b. Males exhibit higher plasma THC concentrations than females. d) Brain THC concentrations measured in rats after the self-administration session shown in a-b. Males exhibit higher brain THC concentrations than females. e) Simple linear regression analyzing the association between the number of infusions earned in the last 30 minutes of THC self-administration (bins 3+4) and plasma THC concentrations. f) Simple linear regression analyzing the association between the number of infusions earned in the last 30 minutes of THC self-administration (bins 3+4) and brain THC concentrations. Both plasma and brain levels positively correlated with the number of infusions earned, r^2^ and p-value in panel. *p<0.05.

Next, analysis of plasma (Fig. 5c) and brain (Fig. 5d) THC levels revealed significant sex differences (plasma: 2way ANOVA main effect of sex F_(1,21)_ = 5.09, p= 0.035; brain: main effect of sex F_(1,19)_ = 4.92, p= 0.039), but no effect of nicotine pre-treatment (plasma p=0.50; brain p=0.74) or interactions (plasma p=0.76; brain p=0.66). Thus, despite overall greater THC self-administration levels in the nicotine pre-treatment group, blood and brain levels of THC were not different based on nicotine exposure. On the other hand, large sex differences in THC levels were observed despite similar amounts of THC self-administration, replicating some reports in the literature ^22,23^, but see ^24^. Given these differences, we questioned if THC concentrations were related to dose self-administered by correlating THC levels to the number of infusions earned in the last two bins, as they were most proximal to blood and tissue collection. Simple linear regression found a significant positive association between infusions earned and plasma (Fig. 5e, p<0.001) and brain (Fig. 5f, p=0.005) levels of THC.

## Discussion

Little is known about the mechanisms mediating differential outcomes from PSU relative to single substance use. Part of this knowledge gap is due to the infrequent use of PSU models in animal research. There are challenges to the design of these experiments as more groups are required to capture the effects of each substance alone and in combination, and in the many ways two different substances can be delivered, such as on different days, combined in the same syringe, or via different routes of administration. Here, we used a concurrent choice dual-catheter model of nicotine and THC use to determine how the availability of both substances affects their self-administration and to determine the effects of nAchR subtype specific antagonists on concurrent self-administration. Similar to our previous findings investigating concurrent choice between nicotine and the synthetic cannabinoid WIN ^21^, we found that nicotine availability increased the number of THC infusions self-administered. However, in the current study the nicotine-enhancement effect during concurrent self-administration was only observed in females, whereas nicotine enhanced WIN self-administration in both sexes. The reason for the difference in males between the two studies is unclear, particularly given that pre-session injection of nicotine did significantly enhance single choice THC self-administration in males. Therefore, nicotine seems to broadly increase propensity for cannabinoid self-administration in both sexes, but there may be subtle cohort effects.

Nicotine has repeatedly been shown to have reinforcement-enhancement effects for otherwise moderate reinforcers, including responding for sensory stimuli like cue lights and sucrose- and cocaine-paired contexts ^25–29^. The ability of nicotine to enhance responding for other drug reinforcers has been less systematically investigated, but studies have found that nicotine pre-treatment increases alcohol and cocaine self-administration ^30–32^, though in one study the ability of nicotine to increase alcohol reinforcement was stronger in females, and can be dose and time dependent ^33^. Nevertheless, the combined use of nicotine with other substances, including cannabis, is likely to contribute to increased acute consumption, which can increase risk for negative outcomes.

In addition, use of multiple substances may lead to differences in neuroplasticity associated with substance use, which may influence the efficacy of therapeutic interventions. Indeed, a prior study investigated the ability of FDA approved treatments for smoking cessation (varenicline) and alcohol use disorder (naltrexone) to reduce ethanol+nicotine intake in female alcohol preferring rats and found that neither treatment reduced consumption of the ethanol+nicotine combination despite being effective for each alone ^34^. Here, we tested whether nAchR antagonists would differentially influence nicotine’s reinforcement enhancing effects, primary nicotine reinforcement, or THC reinforcement in a concurrent choice model and found an interesting pattern of results that differs from prior studies investigating each drug alone. First, THC self-administration was reduced by the β2/4 subunit containing nAchR antagonist, DHβE, but this was only true in females, and appeared to be unrelated to nicotine’s reinforcement enhancing effects, as females with concurrent THC and saline access exhibited a similar reduction. Secondly, nicotine self-administration was not significantly affected by DHβE, unlike prior studies of nicotine alone ^35^, but rather the α7nAchR antagonist, MLA selectively reduced nicotine self-administration in males and females with concurrent access to THC, with no effects in the nicotine+SAL group. Again, this contrasts with prior studies of nicotine self-administration alone that have not found effects of MLA on nicotine’s primary reinforcement or reinforcement enhancing effects ^10,35,36^, but see ^37^. However, the lack of antagonist effects in the nicotine+SAL concurrent access group, also suggests that under choice conditions in general, regardless of what is being self-administered, behavior may be differentially influenced by test agents than when compared to single choice experimental designs. Indeed, when we used a single choice THC self-administration design to more explicitly test nicotine’s reinforcement enhancing effects, without the confound of primary nicotine reinforcement, we observed effects more consistent with the literature where increased THC self-administration in the female nicotine treated group was attenuated by MEC and DHβE, and was unaffected by MLA ^10,12,36^.

Finally, we determined if nicotine exposure affected the bioavailability of THC in the blood or brain. Despite the fact that rats receiving nicotine pre-treatment self-administered more THC than saline controls, and that there were no sex differences in self-administration, the amount of THC measured in blood and brain was significantly higher in males and was not affected by nicotine exposure. Sex differences in THC metabolism have been reported previously ^22^, and our results are consistent with the clinical literature which suggests that blood THC levels are unrelated to behavioral effects ^23^. In addition, these results indicate that nicotine is not causing major changes to blood or brain concentrations of THC to explain the differences in THC self-administration.

Overall, the results of these studies point to important differences in the behavioral effects of drugs when more than one drug is available, and even when rats can respond for two different outcomes, even if one is a saline infusion paired with a discrete cue. Thus, our understanding of the therapeutic value, and even mechanisms of action of some behavioral phenomenon, may be highly dependent on the context in which they are observed. In addition, we found important sex-specific and sex generalizable effects dependent on context, which is also important for interpretation of prior data in the literature. Critically, the results of these studies and others emerging in the literature ^38,39^, point to the importance of incorporating PSU models into investigations of therapeutics for SUDs, and may explain why many promising treatments in animal models have not translated to clinical populations.

## Acknowledgements

The authors would like to thank Victoria Katler and Jennifer Zeak for their technical assistance. The research was supported by USPHS grants P50DA046346, R01DA061227, and R01DA058955, and by operating funds from the Canadian Institute of Health Research (CIHR TCP-431510).

## Statement of Interest

None.

## Data Availability Statement

All data is available upon reasonable request by contacting the corresponding author.

## Notes

### Competing Interest Statement

The authors have declared no competing interest.

